# The species-specificity of energy landscapes for soaring birds, and its consequences for transferring suitability models across species

**DOI:** 10.1101/2021.03.24.436775

**Authors:** Martina Scacco, Eneko Arrondo, J. Antonio Donázar, Andrea Flack, J. Antonio Sánchez-Zapata, Olivier Duriez, Martin Wikelski, Kamran Safi

## Abstract

1. Soaring birds use the energy available in the environment in the form of atmospheric uplifts, to subsidize their flight. Their dependence on soaring opportunities makes them extremely sensitive to anthropogenic wind energy development. Predictive modelling is now considered instrumental to forecast the impact of wind farms on single species of concern. However, as multiple species often coexist in the same area, there is clear need to overcome the limitations of single species approaches.
2. We looked for converging patterns in the way two obligate soaring species use the energy available in the landscape to soar, using movement data from 57 white storks, *Ciconia ciconia*, and 27 griffon vultures, *Gyps fulvus*. We first compared the soaring efficiency of the two species. We then tested the accuracy of topographic features, important correlates of collision risk in soaring birds, in predicting their soaring behaviour. We finally tested the transferability of soaring suitability models across species.
3. Topography alone can predict and map the soaring opportunities available to storks across Europe, but not as efficiently in vultures. Only 20.5% of the study area was suitable to both species to soar, suggesting the existence of species-specific requirements in the use of the landscape for soaring. Storks relied on uplift occurrence while vultures on uplift quality, needing stronger uplifts to support their higher body mass and wing loading.
4. *Synthesis and applications:* Our results indicate that the flight of highly specialized soaring species is more dependent on atmospheric conditions than on static features, and that more knowledge is required to accurately predict their behaviour. Despite the superficially similar soaring behaviour, the two species have different environmental requirements, suggesting that energy landscapes are species-specific. Our models provide a base to explore the effects that changes in the landscape have on the flight behaviour of different soaring species and suggest that there is no reliable and responsible way to shortcut risk assessment in areas where multiple species might be at risk by anthropogenic structures.

## 1 Introduction

The exceptionally high energetic costs of flight selects for minimizing movement costs by taking advantage of the energy provided by the landscape. To flying animals this energy is available in the form of horizontal or vertical air currents (Shepard, Ross & Portugal 2016). Soaring land birds are highly adapted to exploit vertical currents (uplifts) using soaring-gliding flight to a degree that makes them entirely dependent on their availability. By gliding from one uplift to the next, using them as natural lifts, soaring birds can cover large distances with energetic costs shown to be as low as resting level (Duriez et al. 2014). The negligible energetic cost required for soaring-gliding flight is however offset by the disproportionally high cost of flapping flight (Pennycuick 1973, 1972). In fact, the large body mass of soaring birds, combined with proportionately short and broad wings, results in low wing loading (body mass/wing area). This makes them efficient flyers when atmospheric conditions are appropriate, but reduces their manoeuvrability and locomotor efficiency in the absence of uplifts, constraining their movements to areas and times where uplifts are available (Panuccio et al. 2012, Shamoun-Baranes et al. 2017, Vansteelant et al. 2017, Watanabe 2016).

Since the 80s, the increasing demand for reduced emissions of greenhouse gases induces a competition between human infrastructures and flying animals in trying to exploit the energy available in the aerosphere; a competition that Smallwood described as: “(…) wind turbines are simply our means of grabbing energy in which wildlife has already been exploiting for millions of years.” (Köppel 2017). Several studies highlight the high sensitivity of soaring birds to wind farms due to several factors: features in the landscape that generate soaring opportunities are often the same that make wind power plants profitable, raising the rate of encounter of soaring birds wind farms (Nourani & Yamaguchi 2017); limited manoeuvrability and tendency of focusing their attention on the ground while foraging further increases their risk of collision (De Lucas et al. 2008, Marques et al. 2014, Smallwood & Thelander 2008); finally their low annual productivity and slow maturity catalyse low rates of individual losses to irrecoverable population level effects (Allinson 2017, Masden et al. 2010, Smallwood & Thelander 2008, Sanz-Aguilar et al. 2015).

In the last two decades more and more publications are reviewing current measures that aim at mitigating collisions between birds and wind farms while also highlighting the need for further research in this direction (Gove et al. 2013, Laranjeiro et al. 2018, Marques et al. 2014, May et al. 2017, Wang et al. 2015). Planning prior to siting the wind farms is now recognised as first step in the mitigation hierarchy (May et al. 2017), as the siting of both wind farm and specific turbines affect collision risk in birds (Barrios & Rodríguez 2004, Smallwood, Neher & Bell 2009). Predictive modelling, based on previous knowledge about the species’ behaviour in specific environmental contexts, is considered instrumental when it comes to informing investors about the forecasted impact of a particular wind farm on a species of concern. Different modelling studies have shown that topography is one of the primary correlates of increased risk of collision for soaring birds (De Lucas, Ferrer, Bechard & Muñoz 2012, De Lucas et al. 2008, Ferrer et al. 2012, Gove et al. 2013, Watson et al. 2018) due to the soaring opportunities it provides (Sage et al. 2019, Shepard, Williamson & Windsor 2016). Yet, topography has been only rarely considered as an environmental correlate in models predicting fatality rates (De Lucas, Ferrer & Janss 2012, Smallwood, Neher & Bell 2009) or flight behaviour (Aurbach et al. 2018, Becciu et al. 2019, Katzner et al. 2012, Scacco et al. 2019).

We have recently highlighted the role of static topographic features in predicting soaring behaviour and energy expenditure of the white stork *Ciconia ciconia* (Scacco et al. 2019), an obligate soaring bird species. In that study we focused on one single species, like most other studies proposing predictive models of habitat use and collision risk, which are often driven by the urgency of assessing the potential impact on a species of concern (Barrios & Rodríguez 2004, Marques et al. 2014, Smallwood, Rugge & Morrison 2009, Watson et al. 2018). Such focus has clear advantages for the targeted species. Yet, despite the similarities soaring species share in using energy from their environments, and despite their co-occurrence and high susceptibilities to wind farms, different soaring species tend to show different mortality rates caused by collision (De Lucas et al. 2008, Janss 2000, Marques et al. 2014, Martín et al. 2018), highlighting the drawback of focal species analyses. The question arises whether predictive models could be transferred across species boundaries. Comparative studies focusing on the prediction of flight behaviour in specific landscapes allow us to look for common pattern and thus offer an opportunity to generalize and potentially transfer predictive models across species. Such generalized models of flight behaviour can be then complemented with species-specific biologically meaningful variables, and used to predict collision risk in specific areas, in the attempt to maximize mitigation effects at a community level.

In this study we use movement data from two obligate soaring species, the white stork and the griffon vulture *Gyps fulvus*, to search for converging patterns in the way the two species use their energy landscape to soar. Both species are heavily dependent on soaring flight and on the occurrence of uplifts to move across the landscape. We therefore expect them to have similar environmental requirements to sustain their movement. However, they show some morphological differences as well as different foraging strategies: white storks generally forage in open fields and meadows and fly above lowlands; griffon vultures are scavengers, range from lowlands to mountainous landscapes, and have higher body mass and higher wing loading compared to storks (Pennycuick 1972).

In four analytical steps we evaluated to what extent the similar flight behaviour of these two species results in a similar use of the landscape, notwithstanding ecological and morphological differences. 1. As a preparatory step, we used GPS locations and accelerometry data to identify soaring and flapping events of the two species as proxies of low-cost and high-cost flight, respectively (Scacco et al. 2019). 2. We initially considered only the soaring events, and compared the climbing rate (vertical speed) of the two species during soaring, to assess whether differences in their morphology, or in the landscape they are exposed to, affect their soaring efficiency. 3. We then considered both soaring and flapping events, and modelled their occurrence using only topographic features, separately for the two species; these two models were consequently used to predict and map areas potentially suitable for the two species to soar. 4. Finally, we compared suitable areas across species and tested the transferability of our models, that is, if areas suitable to one species could predict the soaring behaviour of the other species.

## 2 Materials and methods

### 2.1 Datasets

We used GPS and tri-axial accelerometry (ACC) data from two obligate soaring species. The dataset included 84 individuals from four different research projects, available on Movebank (Kranstauber et al. 2011): 57 juvenile white storks, tagged in Germany (Flack et al. 2017, 2018), and 27 adult griffon vultures, from two Spanish populations and one French population. The spatial distribution of the dataset defines the extent of the study area (Fig. 1). All animals were equipped with high-resolution, solar GSM-GPS-ACC loggers (e-obs GmbH, Munich, Germany). High-resolution GPS bursts (1 Hz) were collected every 10 or 15 min for 120, 300 or 600 s. ACC data were recorded every 10 min for 3.8 s at 10.54 Hz (40 data points per axis). For details on data collection and data availability see the supplementary material (Table S1).

**Figure 1:**
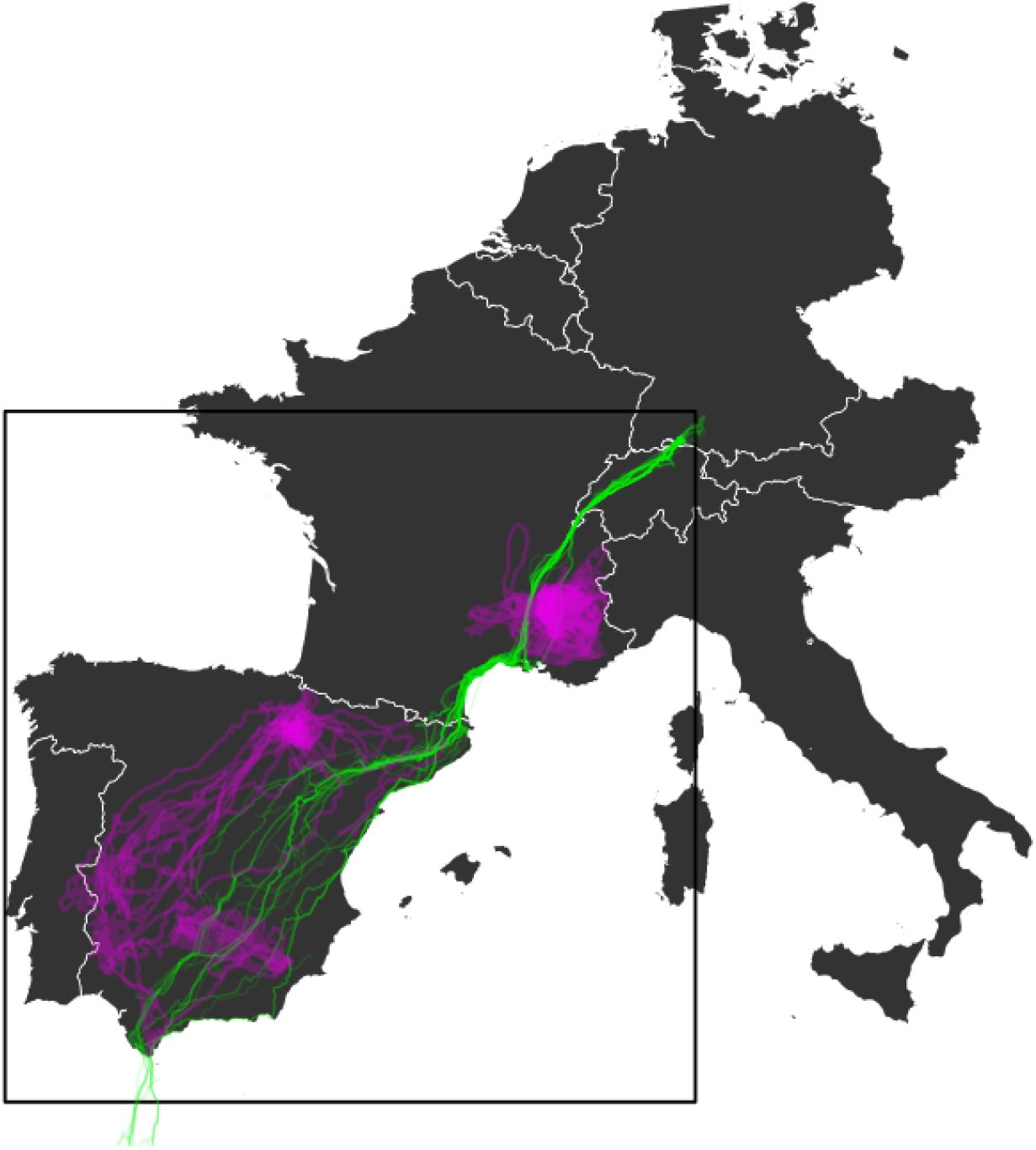
Spatial coverage of the movement data used in the study, white storks in green and griffon vultures in purple. Each line corresponds to the GPS trajectory of an individual bird. The black square indicates the extent of the environmental layers used for the model extrapolation.

### 2.2 Segmentation of the flight behaviour

Soaring events were identified using high-resolution GPS bursts (1 Hz), by applying the Expectation Maximization Binary Clustering on the average vertical speed and the absolute cumulative turning angle calculated on 15 s flight segments (R package `EmbC` (Garriga et al. 2016)). At the end of the procedure, only segments classified as soaring were considered, and the location of each segment was defined by its centroid (mean longitude and latitude). Flapping events were identified using ACC data. We applied k-means clustering with three clusters on DBA-z (Dynamic Body Acceleration on the z-axis) and ODBA (overall DBA) (Wilson et al. 2006)), which have been already used to identify active flight in soaring birds (Duriez et al. 2014, Nathan et al. 2012, Scacco et al. 2019). We finally defined as flapping events the “most active” bursts associated to a heights above ground > 100 m. The location of these events was given by the GPS location closest in time (< 30 s difference). For more details see Scacco et al. (2019) and the supplementary material (S1).

### 2.3 Comparison of vertical speeds

We compared vertical speed of the two species during soaring, while accounting for the different spatial and temporal scales of the two datasets. We modelled vertical speed using a generalized additive model (GAM) including species as categorical predictor, longitude and latitude as interacting thin plate regression splines, and hour of the day as cyclic cubic regression spline. The response variable vertical speed included negative values, therefore we first applied a translation (by adding its minimum value) and then a square-root transformation to meet the assumptions of a Gaussian distribution of the residuals. The model was fit with a Gaussian distribution and run using the R package `mgcv` (Woods 2003).

### 2.4 Soaring suitability models

We modelled the occurrence of soaring and flapping events based on: elevation (digital elevation model, EU-DEM (EEA 2013), terrain unevenness (ruggedness), unevenness in the slope (steepness of a terrain feature), aspect (compass direction faced by a slope) and aspect unevenness (see Hijmans (2016) and supplementary material S3). These topographic variables were included as predictors in a random forest (RF) machine learning algorithm, after verifying the absence of multicollinearity (R package `randomForest` (Liaw & Wiener 2002)). Data from both species were included in two separate models. RF builds many regression trees to distinguish between, and to predict, binary response variables (flapping = 0 *vs* soaring = 1) based on a set of predictor variables. RF is trained with a portion of the data, while the remaining observations (test data) are used to assess model performance. In our case we built a double partitioning: (1) we randomly selected about 80% of the individuals per species (46 storks and 22 vultures), and (2) then applied a 90:10% random partitioning on these individuals and used it to build two RF models (one per species); this second step was repeated 10 times per species, where each time the algorithm was trained with a different 90% of the data and evaluated with the remaining 10%.

The 20% of the individuals left out from the first partitioning did not contribute to the predictive models and were later used to cross-validate the prediction maps extrapolated from the models. Therefore the 90:10% partitioning allows us to measure the performance of the RF in predicting the same pool of individuals the model was built on, whereas the 80:20% partitioning represents a more realistic validation of the extrapolated maps, mimicking the situation in which an external researcher was to use these maps to predict the soaring behaviour of an independent set of individuals (section 2.6).

The performance of the two RF models was evaluated in terms of: (i) area under the curve (AUC) of the receiver operating characteristic (ROC); (ii) sensitivity, proportion of soaring locations correctly classified; (iii) specificity, proportion of flapping locations correctly classified (Franklin 2009). These are threshold-dependent measurements (their values depend on our classification of the predicted probability into 0s and 1s), and were measured at a probability threshold where flapping and soaring were equally well classified (minimum difference between sensitivity and specificity).

### 2.5 Soaring suitability maps

We used the two RF models of soaring suitability (one for each species) and the topographic raster layers corresponding to the RF predictors (spatial granularity of 100 m) to extrapolate two maps of soaring suitability across the study area (Fig. 1). RF, like other machine learning algorithms, is quite unreliable when extrapolating outside the range of the predictors’ values provided for training. We thus omitted raster cells containing environmental values outside that range, and then used each of the ten runs of the RF model to predict the soaring suitability over the manipulated rasters. Each prediction layer (10 per model) was then classified into a binary map (0 or 1) using the threshold where flapping and soaring were classified with the same accuracy (Franklin 2009). We then computed the pixel average of the 10 binary layers, obtaining one final raster per species, with values ranging from 0 to 1. This final prediction map therefore informed us about the soaring suitability in each pixel but also about the model agreement. For the next steps of the analysis we included only pixels with at least 80% agreement, i.e. pixels for which at least 8 out of 10 binary layers agreed on being unsuitable or suitable for soaring (values ≤ 0.2 or ± 0.8). Pixels with values ≤ 0.2 were considered as unsuitable (0), pixels ± 0.8 as suitable (1).

We then compared the soaring suitability maps obtained for the two species and we randomly sampled 100,000 locations (pixels) from areas of the map that were suitable to one or the other species (50,000 per species). We used this dataset to describe and quantify the difference between the two prediction maps in terms of topographic variables used in the models and compared their distribution.

### 2.6 Cross-species prediction of soaring events

For each species we used 20% of the individuals (11 storks and 5 vultures) to test whether the observed soaring events of each species could have been reliably predicted using the soaring suitability map produced from data of either species (in section 2.5). We associated the location of each observed soaring or flapping event to the corresponding soaring suitability value (0 or 1) predicted by both the storks and the griffon vultures’ maps. We then ran two GLMMs (generalized linear mixed effect models) per species, using the observed soaring and flapping events as binary response variable in both models, and using as covariate in one model the soaring suitability predicted by the storks’ map, and in the other model the soaring suitability predicted by the vultures’ map (R package `lme4` (Bates et al. 2014)). Models were fitted using a Bernoulli distribution with a clog-log link function, more appropriate for unbalanced samples (in our case, considerably more soaring than flapping events) (Zuur et al. 2009). Individual identity was included as random intercept in all models. The importance of the predictors was assessed comparing the AIC (Akaike Information Criterion) of each species’ model with the respective null models (one per species, containing only the observed soaring events as response variable and the individual identity as random intercept).

All analyses were run in R (R Core Team 2020).

## 3 Results

### 3.1 Segmentation of the flight behaviour

From the storks’ GPS data, among all individuals, we classified a total of 597.4 hours of flight, of which 311.6 were spent soaring (52% of the recorded flight time). Based on the ACC we could instead classify 102.5 hours of flight, of which only 3.3 hours were classified as flapping (1.9%). From the vultures’ data we classified a total of 1764.4 hours based on the GPS, of which 914.2 spent soaring (51.8%), and 291.6 hours based on the ACC, of which 4.6 classified as flapping (1.6%).

The final dataset, excluding missing topographic information, consisted of 11531 observations for the storks (9797 soaring and 1734 flapping) and 32633 for the vultures (29047 soaring and 3586 flapping).

### 3.2 Comparison of vertical speed

The GAM used to compare the vertical speed of the two species included a total of 37774 soaring events (9180 from storks and 28594 from griffon vultures). Vertical speed was predicted to be 0.33 m/s significantly higher in vultures than in storks [Vultures = 0.1 ± 0.07 (estimate ± s.e.)]. Hour of the day and geographic coordinates had also a significant effect, suggesting that climbing rate would be higher in central hours of the day and in specific regions of the study area (Table S2, Fig. S1).

#### Soaring suitability models

Based on 80% of the individuals (46 storks and 22 vultures) a total of 9901 observations (8488 soaring and 1413 flapping) were available for the stork model and 26673 (24067 soaring and 2606 flapping) for the vultures’ model. The RF model based on stork data resulted in a higher accuracy than the vulture model [AUC stork model: 0.83 ± 0.02; AUC vulture model: 0.71 ± 0.02 (mean ± s.d.)] (Fig. 2). The stork model also resulted in a better ability to discriminate soaring from flapping locations, that is a higher proportion of soaring and flapping locations correctly classified. In fact, the stork model could correctly predict, on average, 75% (± 2%) of the soaring and flapping locations, while only 63% (± 1.5%) were correctly predicted by the vulture model. The complete output of the three models can be found in supplementary material (Table S3). In both the stork and vulture models, the two measures of variable importance (the decrease in accuracy and decrease in node impurity) highlighted terrain elevation, terrain ruggedness and slope unevenness as the most important variables in predicting soaring opportunities. Also the aspect proved to be an important variable, but only in the stork model.

**Figure 2:**
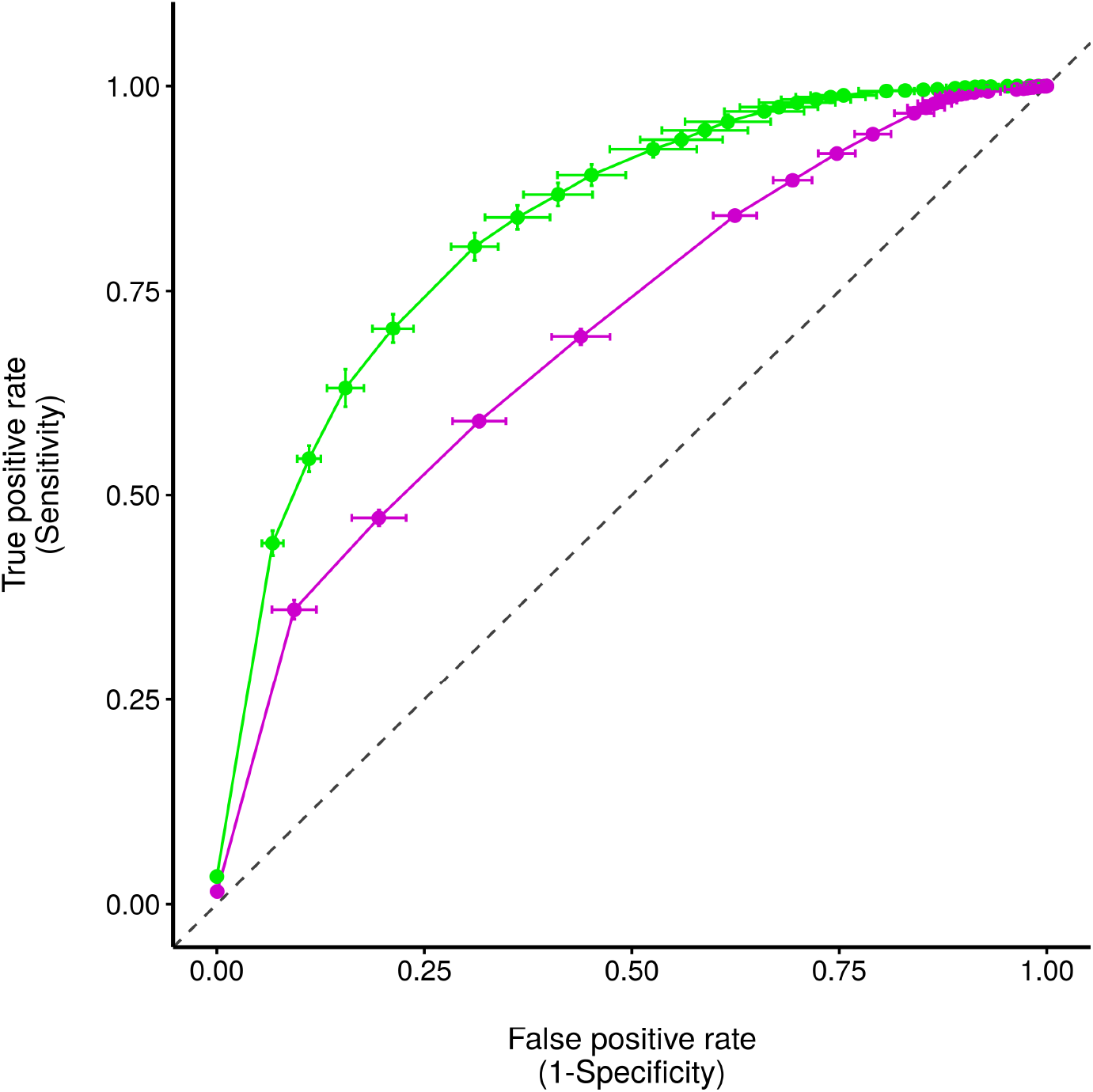
ROC curves of the two soaring suitability models, in green for the storks model and in purple for the vultures model, overlaid for better comparison. The area under the curve (AUC) represents the accuracy of the models, measured on the test dataset. The gray dashed line represents a model whose accuracy is comparable to random (AUC = 0.5). Sensitivity and commission rate values were averaged across the ten runs of each model (solid dots), and the error bars show their standard deviations.

### 3.3 Soaring suitability maps

We used the models to classify an area of about 0.95 million km^2^ for the storks and 0.97 million km^2^ for the vultures (Fig. 3A-B). Only 21% of the storks’ map was predicted to be suitable for them to soar, whereas in the case of vultures’ map, the area suitable to soar was 80%.

**Figure 3:**
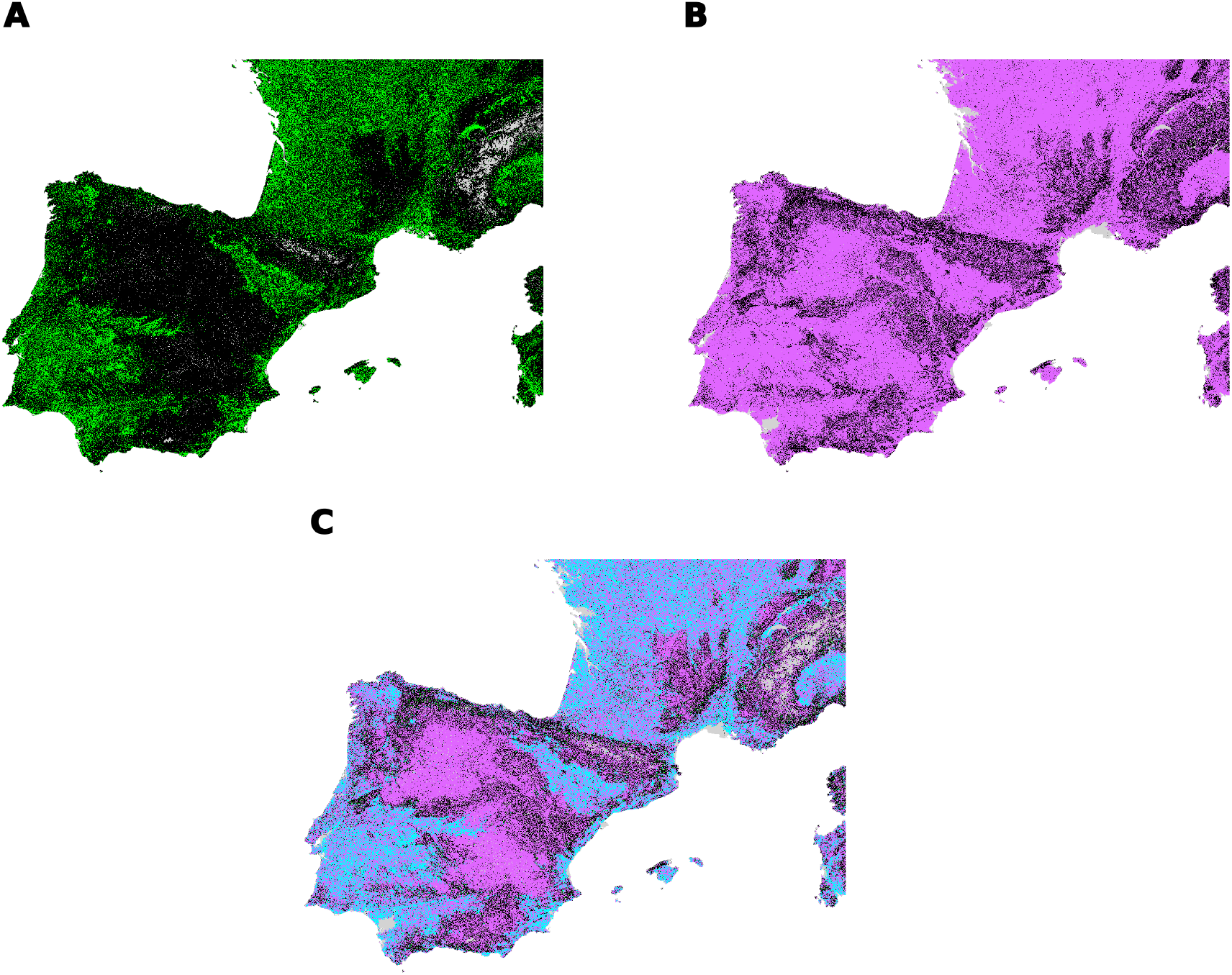
Soaring suitability maps extrapolated from the storks’ (A) and the vultures’ (B) models. Both maps show in colours areas (cells) predicted to be suitable to the species to soar while in black unsuitable cells. Gray represents unclassified cells (containing missing values among the predictors). In (C) a prediction map produced by combining the suitability maps of the two species, showing in light blue cells that are suitable to both species, where the soaring opportunities overlap; in green cells that are available only to storks (not visible due to the small percentage); in purple cells suitable only to vultures; in black cells that are unsuitable to both species; in gray unclassified cells. The three prediction maps will be made available upon acceptance.

We compared the species’ soaring opportunities across 0.86 million km^2^; 20.5% of this area was predicted as suitable to both species, and 18.8% unsuitable to both species to soar (Fig. 3C). 59.9% of the total area was predicted to be suitable to vultures only, whereas only 1.2% was available exclusively to storks, meaning that most of the area suitable to storks was also suitable to vultures, but not vice-versa.

The main difference, in terms of topographic variables, between areas suitable to one or the other species, concerned the aspect. Storks, unlike vultures, showed to be selective in terms of aspect, with a peak in the distribution around 200 degrees (S-SW). The distribution of the other topographic variables did not highlight any other species-specific difference (Fig. 4).

**Figure 4:**
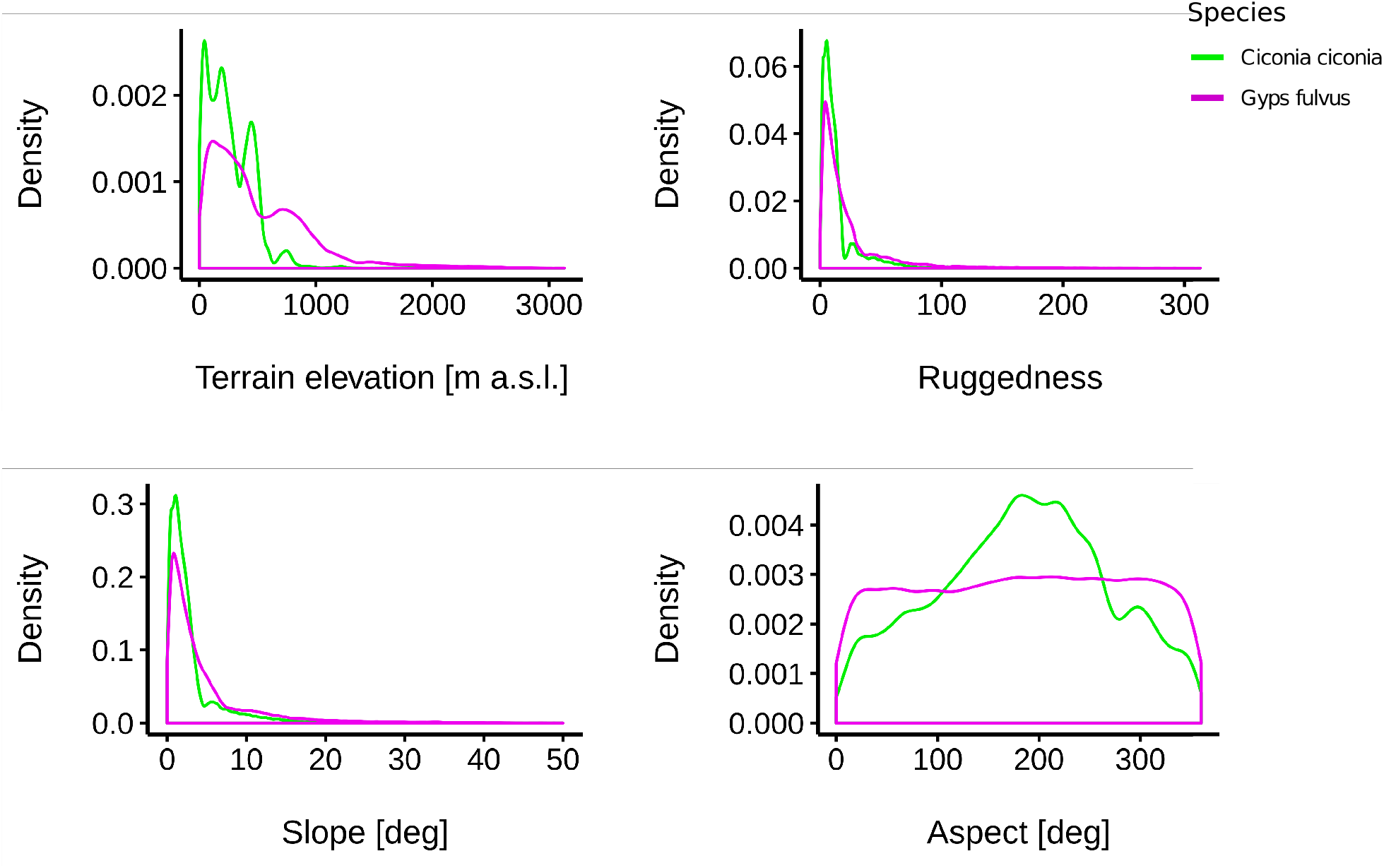
Density distribution of four topographic variables (terrain elevation, ruggedness, slope unevenness and aspect), extracted from 50,000 cells per species, randomly selected from areas predicted to be suitable to the storks and to the vultures to soar.

### 3.4 Cross-species prediction of soaring events

The two stork models (one using the storks’ map as covariate, one using the vultures’ map) showed that the probability of soaring significantly increased in areas predicted as suitable by either species, but was stronger for the model including the prediction based on storks [Storks’ suitability map = 1.42 ± 0.12; Vultures’ suitability map = 0.59 ± 0.15 (estimate ± s.e.)]. The AIC of both of the stork models was lower compared to the respective null model [AIC storks’ suitability map = 695.8; vultures’ suitability map = 859.5; null model = 870.71], suggesting that adding a static soaring suitability map as covariate helps us predicting their soaring behaviour (Table 1A).

**Table 1:**
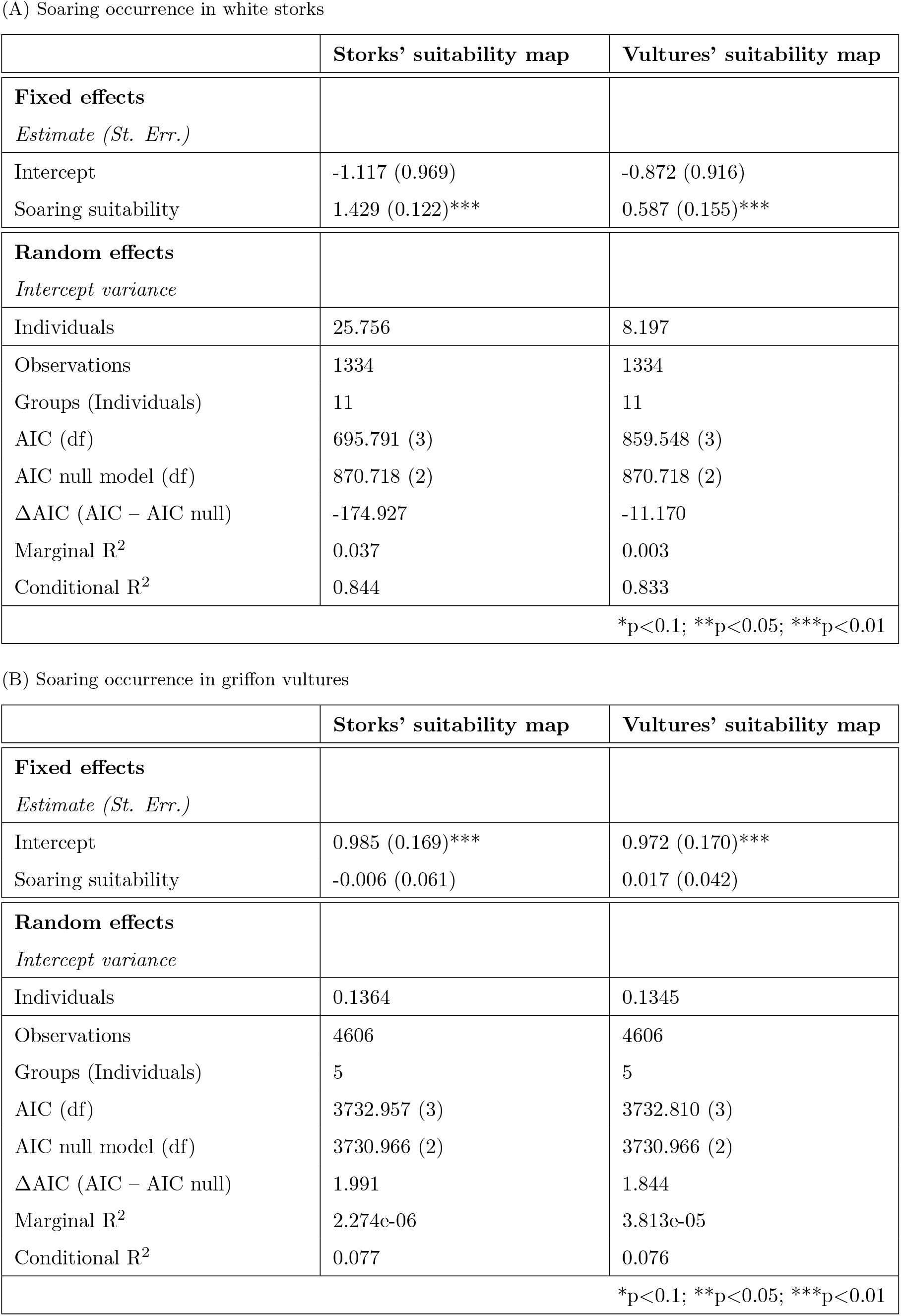
Output of the GLMMs explaining the observed soaring probability of the two species as a function of the soaring suitability predicted by the models of each species.

(A) Soaring occurrence in white storks
|  | Storks' suitability map | Vultures' suitability map |
| --- | --- | --- |
| **Fixed effects** |  |  |
| *Estimate (St. Err.)* |  |  |
| Intercept | -1.117 (0.969) | -0.872 (0.916) |
| Soaring suitability | 1.429 (0.122)*** | 0.587 (0.155)*** |
| **Random effects** |  |  |
| *Intercept variance* |  |  |
| Individuals | 25.756 | 8.197 |
| Observations | 1334 | 1334 |
| Groups (Individuals) | 11 | 11 |
| AIC (df) | 695.791 (3) | 859.548 (3) |
| AIC null model (df) | 870.718 (2) | 870.718 (2) |
| ΔAIC (AIC – AIC null) | -174.927 | -11.170 |
| Marginal R<sup>2</sup> | 0.037 | 0.003 |
| Conditional R<sup>2</sup> | 0.844 | 0.833 |
| *p<0.1; **p<0.05; ***p<0.01 |  |  |

(B) Soaring occurrence in griffon vultures
|  | Storks' suitability map | Vultures' suitability map |
| --- | --- | --- |
| **Fixed effects** |  |  |
| *Estimate (St. Err.)* |  |  |
| Intercept | 0.985 (0.169)*** | 0.972 (0.170)*** |
| Soaring suitability | -0.006 (0.061) | 0.017 (0.042) |
| **Random effects** |  |  |
| *Intercept variance* |  |  |
| Individuals | 0.1364 | 0.1345 |
| Observations | 4606 | 4606 |
| Groups (Individuals) | 5 | 5 |
| AIC (df) | 3732.957 (3) | 3732.810 (3) |
| AIC null model (df) | 3730.966 (2) | 3730.966 (2) |
| ΔAIC (AIC – AIC null) | 1.991 | 1.844 |
| Marginal R<sup>2</sup> | 2.274e-06 | 3.813e-05 |
| Conditional R<sup>2</sup> | 0.077 | 0.076 |
| *p<0.1; **p<0.05; ***p<0.01 |  |  |

In contrast, both vulture models showed a weak and non significant relationship between the soaring suitability values predicted by the two maps and the observed soaring events. The effect size was small in both cases [Storks’ suitability map = −0.005 ± 0.06; Vultures’ suitability map = 0.016 ± 0.04 (estimate ± s.e.)] and the AIC of both models was comparable to the null model [AIC storks’ suitability map = 3732.9; vultures’ suitability map = 3732.8; null model = 3730.9]. Consequently and in contrast to the stork models, both the static soaring suitability maps did not contribute significantly to improve predicting the soaring behaviour of vultures (Table 1B).

## 4 Discussion

Predictive modelling of flight behaviour on single species is instrumental to inform the siting of anthropogenic infrastructures and to minimize collision risk. However, multiple species of concern often coexist in the same area, and our comparative study is a first attempt to test the transferability of predictive models across species with similar flight behaviour.

Our results indicate that, despite the superficially similar soaring behaviour, white storks and griffon vultures have different environmental requirements and that soaring suitability models cannot and should not be transferred between species. The soaring suitability maps extrapolated from our models showed that only 20.5% of the classified study area was available to both species to soar. Most of the area suitable to the storks was also available to the vultures, but not vice-versa, implying that vultures had a larger area potentially available for soaring. Vultures had a higher vertical speed, soared in a variety of landscape conditions and contrary to storks, their soaring flight was not related to a specific range of aspect values. Finally, our results concerning model transferability showed that a model based on one species performed poorly in predicting the soaring events of the second species.

Both the larger area available to soar and the less specific topographic requirements, might depict vultures as more flexible fliers. Their vertical speed was higher than in storks, which is surprising considering their higher body mass and higher wing loading (Pennycuick 1972). Due to their morphology, vultures are expected to need stronger uplifts to fly efficiently and to depend more on the specific environmental conditions able to produce such support (Pennycuick 1973, Shamoun-Baranes et al. 2003). Our interpretation of their higher climbing rate is thus that they used disproportionally stronger uplifts (better in quality) than those used on average by storks. This suggests that storks are able to take advantage of weaker uplifts compared to vultures, and it seems enough for them to rely on their occurrence rather than on their quality. In contrast, vultures need stronger uplifts to support their larger mass; they therefore rely on uplift quality and should be more selective in the soaring conditions.

Our results seem thus contradictory: they suggest that vultures should be using stronger uplifts, which can only be generated under specific environmental conditions, but they depict vultures as flexible fliers, having most of the study area potentially available to soar. We believe that the source of this contradiction is that our soaring suitability models are based exclusively on static topography-related covariates. In a recent study we showed that static topographic variables are effective in predicting the occurrence of uplifts used by storks. However the same variables were not useful predictors of their climbing rate in thermals (Scacco et al. 2019), for which the use of weather covariates is expected to be crucial (Aurbach et al. 2018, Becciu et al. 2019). In this follow up study, our static model based on topography confirmed its high accuracy in predicting soaring occurrence in storks, but had a lower accuracy in vultures, both in the train and test datasets. This suggests that topography alone cannot predict vultures’ soaring behaviour, probably due to lacking predictive performance in uplift quality, which they heavily rely on. Consequently the contradiction emerged from our results represent shortcomings of the methods used to predict the occurrence of soaring behaviour in vultures.

This interpretation has implications also on modelling species occurrence at global scale. The higher the wing loading of a species, the more its movement and therefore distribution will be restricted in space and time not only to where, but mainly when, uplift conditions are optimal to provide enough lift (Williams et al. 2020). The flight of such specialized species will therefore be more dependent on atmospheric conditions than on static features, which implies that more knowledge is required to accurately predict it (Soultan & Safi 2017).

The link between species’ movement capacity, biogeography and species distribution models is often overlooked, or when taken into account, it mostly relates to the obstacle posed by large ecological barriers to colonization processes (Cumming et al. 2012, Mellone 2020). In highly specialised soaring species, considering the link between biogeography and movement only in relation to colonization would underestimate the role of uplifts as essential part of the niche of these species, for which uplifts should be considered among the suite of resources they need in order to exist in a certain area. Long accepted definitions of ecological barriers such as water bodies for soaring birds, are now being revisited as more flexible than previously thought (Nourani et al. 2020), and should be adapted to accommodate the diversity in movement capacity among species. The aim of this study was to predict the energetic value of different areas of the landscape, distinguishing areas providing soaring opportunities from those that did not; thus, we did not further differentiate between types of soaring or types of uplift. Yet, different species are known to rely on different types of uplifts and we recognise the use of thermal versus orographic uplifts as being another important factor potentially influencing the predictability of flight behaviour, given the different mechanisms that generate them (Duerr et al. 2014, Poessel et al. 2018).

Our predictive models of soaring opportunities represent an attempt to describe the potential energy available in the landscape to allow soaring species to move efficiently, and can be used as base layers in movement ecology analyses. The distribution of soaring opportunities is one of the resources needed by these species to exist in an area. However, species-specific patterns of use of the landscape are also mediated by other biologically relevant factors, not targeted by this study, such as the distribution of food resources or nesting opportunities. These factors vary extensively among species and have been found to be related to vulture fatalities at wind farms (Carrete et al. 2012). These are therefore essential variables to consider before using these models to inform the planning of anthropogenic infrastructures.

## 5 Conclusions

Static landscape structure greatly influences the energy available to soaring birds in the landscape and thus their movement, but for some species static variables are not sufficient alone and need to be associated to weather information to produce more reliable predictions of flight behaviour. We suggest that using the soaring behaviour of the species to detect uplift events is instrumental to isolate those uplifts that can be effectively detected and used by that species to fly. However, once we use an environmental model to extrapolate the occurrence of those uplifts over large areas, it becomes really tempting to apply this same prediction to species having a similar flight behaviour. The results of this study warn against this practice and clarify that such maps represent the energy available in the landscape from the point of view of the species used to build the model.

Energy landscapes are therefore species-specific, meaning that the same landscape varies in the soaring opportunities it offers to different species, and affects their movement pattern differently. The use of species-specific prediction models would allow us for more flexibility in the choice of the important variables to consider, including meaningful biological information that differ among species. Accounting for these differences is crucial when predicting movement and landscape connectivity and there seems to be no reliable and responsible way to shortcut risk assessment in areas where multiple species might be at risk by anthropogenic structures.

## Authors’ contributions

MS KS and MW conceived and designed the study, EA JAD AF JASZ and OD provided the tracking data, MS and KS carried out the analyses, interpreted the results and wrote the first draft of the manuscript. All authors contributed suggestions and text to subsequent drafts. All authors gave final approval for publication.

## Acknowledgements

We acknowledge funding from the Max Planck Institute of Animal Behavior. MS was supported by the German Academic Exchange Service (DAAD) and by the International Max Planck Research School for Organismal Biology. Environmental products courtesy of European Environmental Agency. The collection of the Spanish vultures’ data was funded by the Project RNM-1925 (Junta de Andalucía) and the Comunidad de Bardenas Reales de Navarra.

