## Supplementary material for "The species-specificity of energy landscapes for soaring birds, and its consequences for transferring suitability models across species"

**S4 Cross-species prediction of soaring events** 11

### S1 Dataset and segmentation

#### S1.1 Dataset

The GPS and tri-axial accelerometry data used in this study were from two species and collected in four different research projects, available on Movebank (Kranstauber et al. 2011) (Fig. 1 in main text). All animals included in the study were equipped with high-resolution, solar GSM-GPS-ACC loggers (e-obs GmbH, Munich, Germany).

The white storks' dataset included 61 immature white storks (*Ciconia ciconia*) during their first migration. Data were collected by Max Planck of Animal Behavior (Flack et al. 2017, Weinzierl et al. 2016) and were deposited in the Movebank Data Repository (`http://dx.doi.org/10.5441/001/1.bj96m274`). Loggers recorded one GPS location every 5 min between 2:00 and 20:00 GMT. If instantaneous ground speed was > 2 m/s, bursts of high-resolution GPS locations (1 Hz) were recorded every 15 min for 120 or 300 s. In addition to the GPS locations, tri-axial accelerometry (ACC) was recorded every 10 min for a duration of 3.8 s at a sampling rate of 10.54 Hz (40 data points per axis). High-resolution GPS recordings were collected from August to September 2014.

The vultures' dataset originally included a total of 75 adult griffon vultures (*Gyps fulvus*) from two populations in Spain (Ebro valley in northern Spain and Cazorla in southern Spain) and one population in the French Alps (Baronnies). Data from the Spanish populations were all collected by the VultureGroup, from December 2014 to date, and included 37 and 31 individuals (Movebank study names: "Griffon vulture - Bardenas Reales" and "Scavengers and wild ungulates", respectively). High-resolution GPS recordings associated with ACC were available for 20 individuals between April and July 2019, with the same sampling schedule used for the white storks. Data from the French population were collected by the Centre d'Ecologie Fonctionnelle et Evolutive and Vautours en Baronnies, from January 2015 to date (Movebank study name: "Eurasian Griffon Vulture in France (Alps - Baronnies)"). The sampling schedule of these tags was the same for the ACC but slightly different for the GPS: one GPS location every 1 min between 7:00 and 17:00 GMT; high resolution GPS bursts (1 Hz) were recorded every 10 min for 300 or 600 s. The high resolution GPS bursts were available for 7 individuals for the entire period of data collection. See Table S1 for an overview.

Table S1: Overview of data collection (sampling schedule) and data availability [HR = high resolution 1 Hz GPS].

| Species | Population | N. indiv. | N. HR indiv. | Age | HR tracking period | GPS | ACC | Data owner | Movebank data source |
|---|---|---|---|---|---|---|---|---|---|
| White storks (*Ciconia ciconia*) | Germany | 61 | 57 | Immatures, first migration | August - September 2014 | 1 position every 15 min + bursts at 1 Hz every 15 min for 120 s or 300 s | Bursts every 10 min at 10.54 Hz for 3.8 s | MPI of Animal Behavior | `http://dx.doi.org/10.5441/001/1.bj96m274` (Flack et al. 2017) |
| Griffon vultures (*Gyps fulvus*) | Spain - Ebro valley | 37 | 12 | Adult, resident | April - July 2019 | As for storks | As for storks | Vulture Group | Griffon vulture - Bardenas Reales |
| | Spain - Cazorla | 31 | 8 | Adults, resident | April - July 2019 | As for storks | As for storks | Vulture Group | Scavengers and wild ungulates |
| | France Baronnies | 7 | 7 | Adults, resident | January 2015 to date | 1 position every 1 min + bursts at 1 Hz every 10 min for 300 s or 600 s | As for storks | Centre d'Ecologie Fonctionnelle et Evolutive | Eurasian Griffon Vulture in France (Alps - Baronnies) |

##### S1.2 Segmentation

Soaring events were identified based on high-resolution GPS bursts with a duration of at least 120 s. We first grouped the 1 Hz locations in track segments of 15 s (average duration of one complete soaring circle (Weinzierl et al. 2016)). We then applied the Expectation Maximization Binary Clustering on the average vertical speed and the absolute cumulative turning angle calculated on these segments (R package `EmbC` (Garriga et al. 2016)). At the end of the procedure, each 15 s segment along the animal trajectory was individually assigned to one of three behavioural categories: circular soaring, linear soaring and gliding. The purpose of the subsequent analysis was to distinguish between the use of active versus passive flight; we therefore excluded gliding segments and assigned linear and circular soaring to the same category. The location of each soaring segment was defined by its centroid (mean longitude and latitude). For more details see Scacco et al. (2019).

Flapping events were identified based on ACC data using DBA-z (Dynamic Body Acceleration on the z-axis) and ODBA (overall DBA), which have been shown to be good proxies for energy expenditure and have already been used to identify active flight in soaring birds (Duriez et al. 2014, Nathan et al. 2012, Scacco et al. 2019). We first calculated ODBA and DBA-z for each ACC burst following Wilson et al. (2006); and then applied k-means clustering to group the ACC bursts into three classes based on the amount of activity recorded: least active, intermediately active, and most active. We finally defined as flapping events the "most active" bursts associated to a heights above ground $> 100$ m. The location of these events was given by the GPS location closest in time (less than 30 s difference). For more details on the segmentation procedure see Scacco et al. (2019).

#### S2 Comparison of climbing rate

We compared the climbing rate of the two species during soaring, while accounting for the different spatial and temporal scales of the two datasets. We modelled climbing rate using a generalized additive model (GAM) including species as categorical predictor, longitude and latitude as interacting thin plate regression splines, and hour of the day as cyclic cubic regression spline. The response variable climbing rate included negative values, therefore we first applied a translation (by adding its minimum value) and then a square-root transformation to meet the assumptions of a Gaussian distribution of the residuals. The complete model output is shown in Table S2. See also Fig. **??** for a visualization of the partial effect of some predictors on the species' climbing rates.

Table S2: Summary of the climbing rate model (GAM output).

| | **Response: sqrt(climbing rate)** | |
|---|---|---|
| **Fixed effects** | **Estimate (St. Err.)** | **t value** |
| Intercept | 1.710 (0.005) | 318.156*** |
| Species *Gyps fulvus* | 0.095 (0.007) | 13.643*** |
| **Smooth terms** | **edf** | **F** |
| $s$(Longitude, Latitude) | 25.756 | 18.06*** |
| $s$(Hour of the day) | 5.987 | 153.25*** |
| Observations | 37774 | |
| Adjusted $R^2$ | 0.099 | |
| | | *p<0.1; **p<0.05; ***p<0.01 |

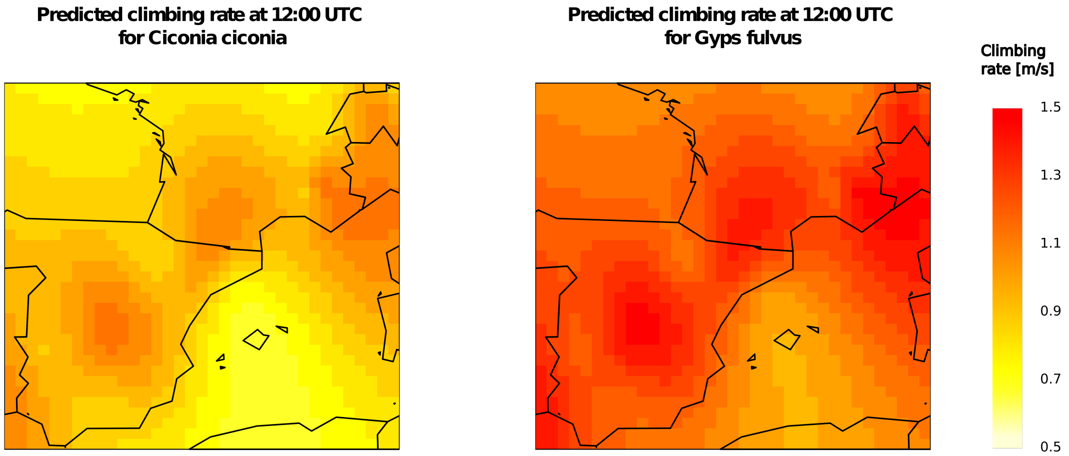

(A)

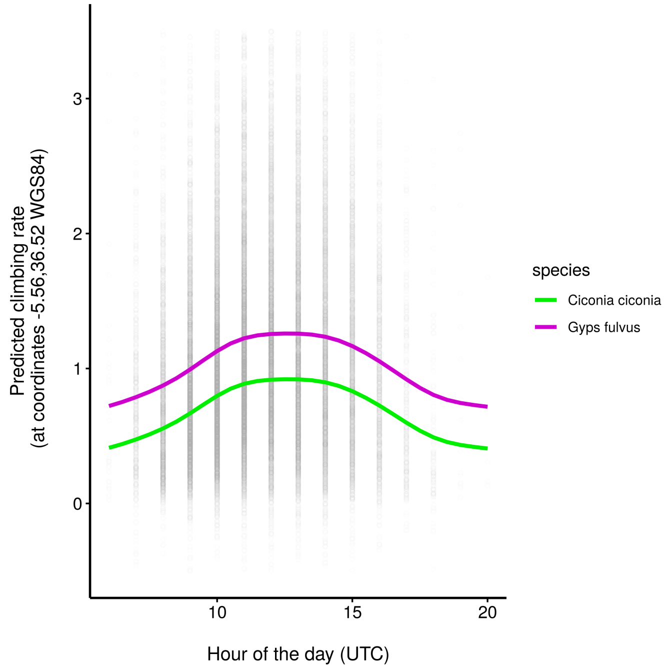

(B)

Figure S1: Non linear relationship between two predictors and the response variable climbing rate. In (A) the effect of spatial location (longitude/latitude) on the climbing rate of storks and vultures at a fixed time (12:00 UTC), with lighter colours indicating locations that predicted a higher climbing rate. In (B) the effect of hour of the day at a fixed location.

### S3 Soaring suitability

#### S3.1 Environmental variables

In order to characterize the surface features we used the publicly available elevation map EU-DEM (based on SRTM and ASTER Global Digital Elevation Model) (EEA 2013)and we computed slope, aspect and ruggedness (topographic heterogeneity) using the R package `raster` (Hijmans 2016). The native spatial granularity of the elevation map is 1 arcsec (about 25 m near the Equator). Raster cells were aggregated to 100 m resolution. Slope and aspect were computed according to Horn (Horn 1981). The ruggedness was calculated as the difference between the maximum and the minimum value of a cell and its 8 surrounding cells. Unevenness in the aspect, the slope and the elevation (this last one also called Topographic Position Index) were computed as the difference between the value of a cell and the mean value of its 8 surrounding cells. Highly correlated layers were excluded from the model to avoid multicollinearity (this was the case of the slope because of the high correlation with the ruggedness).

#### S3.2 Soaring suitability models

Table S3: Random forest evaluation of the two soaring suitability models, based on the test set and averaged across the ten cross-validations. (A) Storks' model AUC: 0.83 ± 0.02 (mean ± s.d.); (B) Vultures' model AUC : 0.71 ± 0.02.

(A)

| **Threshold** | **Sensitivity** | **Specificity** | **Commission Error** | **Omission Error** | **TSS** |
|---|---|---|---|---|---|
| 0 | 1 (0) | 0 (0) | 1 (0) | 0 (0) | 0 (0) |
| 0.05 | 1 (0) | 0.021 (0.011) | 0.979 (0.011) | 0 (0) | 0.021 (0.011) |
| 0.1 | 1 (0) | 0.035 (0.012) | 0.965 (0.012) | 0 (0) | 0.035 (0.012) |
| 0.15 | 1 (0) | 0.056 (0.015) | 0.944 (0.015) | 0 (0) | 0.056 (0.015) |
| 0.2 | 0.999 (0.001) | 0.078 (0.02) | 0.922 (0.02) | 0.001 (0.001) | 0.077 (0.021) |
| 0.25 | 0.998 (0.002) | 0.098 (0.023) | 0.902 (0.023) | 0.002 (0.002) | 0.097 (0.023) |
| 0.3 | 0.997 (0.002) | 0.118 (0.025) | 0.882 (0.025) | 0.003 (0.002) | 0.115 (0.025) |
| 0.35 | 0.996 (0.002) | 0.149 (0.022) | 0.851 (0.022) | 0.004 (0.002) | 0.144 (0.022) |
| 0.4 | 0.994 (0.003) | 0.193 (0.034) | 0.807 (0.034) | 0.006 (0.003) | 0.187 (0.033) |
| 0.45 | 0.989 (0.004) | 0.245 (0.04) | 0.755 (0.04) | 0.011 (0.004) | 0.234 (0.039) |
| 0.5 | 0.984 (0.006) | 0.278 (0.042) | 0.722 (0.042) | 0.016 (0.006) | 0.262 (0.04) |
| 0.55 | 0.974 (0.007) | 0.323 (0.047) | 0.677 (0.047) | 0.026 (0.007) | 0.297 (0.045) |
| 0.6 | 0.963 (0.008) | 0.362 (0.059) | 0.638 (0.059) | 0.037 (0.008) | 0.325 (0.058) |
| 0.65 | 0.946 (0.009) | 0.412 (0.052) | 0.588 (0.052) | 0.054 (0.009) | 0.358 (0.054) |
| 0.7 | 0.923 (0.01) | 0.474 (0.052) | 0.526 (0.052) | 0.077 (0.01) | 0.397 (0.055) |
| 0.75 | 0.891 (0.013) | 0.548 (0.041) | 0.452 (0.041) | 0.109 (0.013) | 0.44 (0.049) |
| 0.8 | 0.84 (0.015) | 0.638 (0.039) | 0.362 (0.039) | 0.16 (0.015) | 0.478 (0.044) |
| 0.85 | 0.758 (0.016) | 0.751 (0.027) | 0.249 (0.027) | 0.242 (0.016) | 0.509 (0.035) |
| 0.9 | 0.631 (0.023) | 0.845 (0.022) | 0.155 (0.022) | 0.369 (0.023) | 0.476 (0.035) |
| 0.95 | 0.441 (0.015) | 0.933 (0.013) | 0.067 (0.013) | 0.559 (0.015) | 0.374 (0.019) |
| 1 | 0.034 (0.006) | 1 (0) | 0 (0) | 0.966 (0.006) | 0.034 (0.006) |

(B)

| **Threshold** | **Sensitivity** | **Specificity** | **Commission Error** | **Omission Error** | **TSS** |
|---|---|---|---|---|---|
| 0 | 1 (0) | 0 (0) | 1 (0) | 0 (0) | 0 (0) |
| 0.05 | 1 (0) | 0 (0) | 1 (0) | 0 (0) | 0 (0) |
| 0.1 | 1 (0) | 0.001 (0.002) | 0.999 (0.002) | 0 (0) | 0.001 (0.002) |
| 0.15 | 1 (0) | 0.006 (0.005) | 0.994 (0.005) | 0 (0) | 0.006 (0.006) |
| 0.2 | 0.999 (0.001) | 0.011 (0.007) | 0.989 (0.007) | 0.001 (0.001) | 0.01 (0.007) |
| 0.25 | 0.999 (0.001) | 0.016 (0.011) | 0.984 (0.011) | 0.001 (0.001) | 0.015 (0.012) |
| 0.3 | 0.998 (0.001) | 0.018 (0.012) | 0.982 (0.012) | 0.002 (0.001) | 0.016 (0.012) |
| 0.35 | 0.998 (0.001) | 0.024 (0.012) | 0.976 (0.012) | 0.002 (0.001) | 0.021 (0.012) |
| 0.4 | 0.996 (0.001) | 0.036 (0.014) | 0.964 (0.014) | 0.004 (0.001) | 0.032 (0.014) |
| 0.45 | 0.993 (0.002) | 0.071 (0.015) | 0.929 (0.015) | 0.007 (0.002) | 0.064 (0.016) |
| 0.5 | 0.991 (0.003) | 0.098 (0.019) | 0.902 (0.019) | 0.009 (0.003) | 0.089 (0.02) |
| 0.55 | 0.989 (0.003) | 0.11 (0.018) | 0.89 (0.018) | 0.011 (0.003) | 0.099 (0.02) |
| 0.6 | 0.985 (0.003) | 0.122 (0.021) | 0.878 (0.021) | 0.015 (0.003) | 0.107 (0.022) |
| 0.65 | 0.979 (0.004) | 0.135 (0.02) | 0.865 (0.02) | 0.021 (0.004) | 0.114 (0.022) |
| 0.7 | 0.967 (0.004) | 0.16 (0.024) | 0.84 (0.024) | 0.033 (0.004) | 0.126 (0.024) |
| 0.75 | 0.941 (0.004) | 0.21 (0.022) | 0.79 (0.022) | 0.059 (0.004) | 0.151 (0.023) |
| 0.8 | 0.885 (0.004) | 0.306 (0.023) | 0.694 (0.023) | 0.115 (0.004) | 0.191 (0.025) |
| 0.85 | 0.778 (0.004) | 0.459 (0.038) | 0.541 (0.038) | 0.222 (0.004) | 0.236 (0.042) |
| 0.9 | 0.59 (0.007) | 0.684 (0.032) | 0.316 (0.032) | 0.41 (0.007) | 0.274 (0.031) |
| 0.95 | 0.36 (0.012) | 0.907 (0.027) | 0.093 (0.027) | 0.64 (0.012) | 0.267 (0.025) |
| 1 | 0.015 (0.003) | 1 (0.001) | 0 (0.001) | 0.985 (0.003) | 0.015 (0.003) |

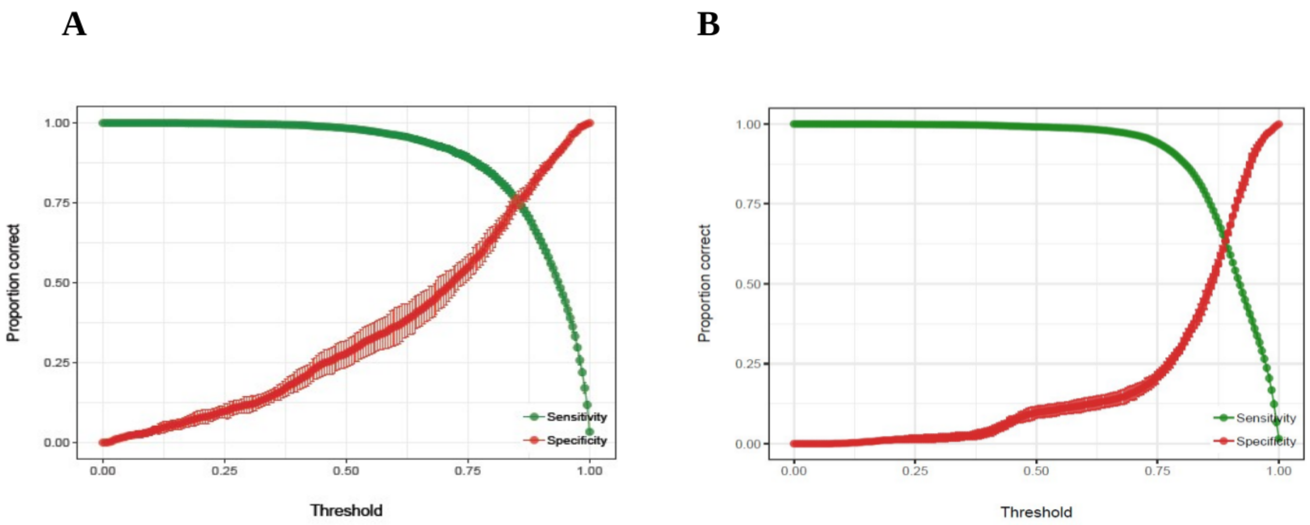

Figure S2: Accuracy of the two uplift suitability models, storks (A) and vultures (B), in terms of sensitivity (proportion of soaring locations correctly classified, in green) and specificity (proportion of flapping locations correctly classified, in red) at different thresholds values. The solid points represent the value of Sensitivity and Specificity, averaged across the ten runs of each model, at a threshold that maximize the value of the True Skill Statistics. The error bars show their standard deviations.

#### S4 Cross-species prediction of soaring events

A

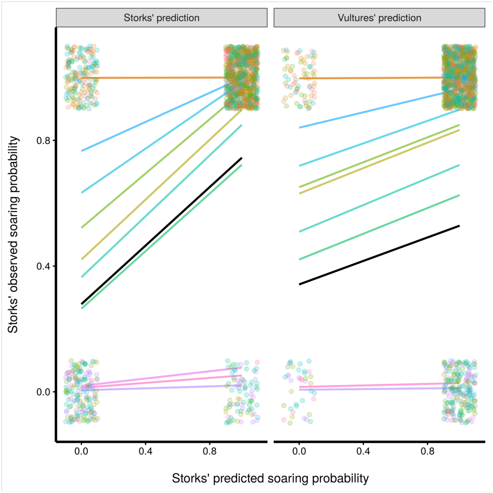

B

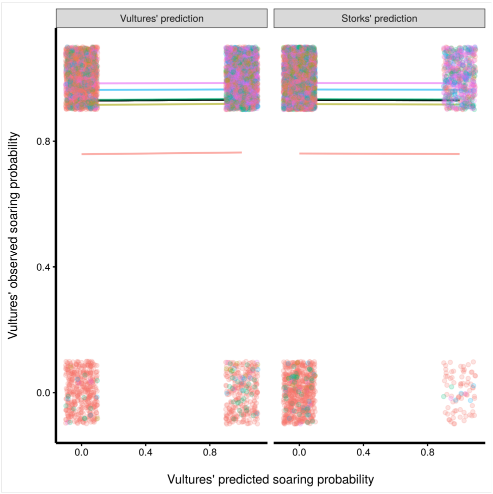

Figure S3: Accuracy of the two uplift suitability models, storks (A) and vultures (B), in terms of sensitivity (proportion of soaring locations correctly classified, in green) and specificity (proportion of flapping locations correctly classified, in red) at different thresholds values. The solid points represent the value of Sensitivity and Specificity, averaged across the ten runs of each model, at a threshold that maximize the value of the True Skill Statistics. The error bars show their standard deviations.
